# High *GPX4* Expression Marks an Oxidative-Metabolic, PD-L1/CD47-Low and Stroma-Poor Tumor State in Hepatocellular Carcinoma

**DOI:** 10.64898/2026.11.13.766360

**Authors:** Prakash Raj Govindaraj, Madu Pascal Akaye

**Affiliations:** Moscow Institute of Physics and Technology, Moscow, Russia; Institute of Personalized Oncology, Sechenov First Moscow State Medical University, Moscow, Russia

**Keywords:** hepatocellular carcinoma, ferroptosis, GPX4, PD-L1, CD47, tumor microenvironment, NRF2, gene set enrichment analysis, TCGA

## Abstract

**Background:** Glutathione peroxidase 4 (*GPX4*) protects cancer cells from ferroptosis and is a candidate therapeutic target in hepatocellular carcinoma (HCC). How *GPX4* expression relates to the immune and metabolic features of patient tumors is not well characterized.

**Methods:** We analyzed RNA-sequencing data from 366 primary tumors of the TCGA-LIHC PanCancer Atlas cohort (365 with survival data). Associations between *GPX4* and ten immune and stress-response markers were tested by Spearman correlation with Benjamini-Hochberg correction, then re-estimated after adjustment for consensus tumor purity, *CTNNB1* and *TP53* mutation, age and sex. A T-cell-inflamed profile, a CD8 T-cell score and an NRF2 target-gene score, gene set enrichment analysis (GSEA) of *GPX4*-high versus *GPX4*-low tumors, and Cox models were added as further tests.

**Results:** Higher *GPX4* expression was associated with lower expression of PD-L1 (*CD274*; ρ = −0.34, 95% CI −0.43 to −0.24, p = 2.4 × 10^−11^), *CD47* (ρ = −0.21), the macrophage marker *CD163* (ρ = −0.17) and a fibroblast/extracellular-matrix score (ρ = −0.22). These associations persisted after adjustment (*CD274* ρ = −0.37 in the fully adjusted model). GSEA showed enrichment of oxidative phosphorylation, mitochondrial, fatty-acid, xenobiotic and glutathione metabolism programs and depletion of extracellular-matrix, TGF-β and cell-adhesion programs in *GPX4*-high tumors. *GPX4* was not associated with a T-cell-inflamed profile (ρ = −0.07), a CD8 T-cell score (ρ = −0.03), cytolytic activity (ρ = 0.04) or an NRF2 target-gene score (ρ = 0.03), although *NFE2L2* mRNA was inversely correlated with *GPX4* (ρ = −0.32). *GPX4* expression was not associated with overall survival (stage-adjusted hazard ratio per SD 1.09, 95% CI 0.91-1.29).

**Conclusions:** In TCGA-LIHC, high *GPX4* expression marks a metabolically active, PD-L1/*CD47*-low and matrix-poor tumor state, with no evidence of T-cell exclusion, NRF2 pathway activation or prognostic value. These correlative findings come from a single cohort and require external validation and functional testing.

## INTRODUCTION

Hepatocellular carcinoma (HCC) is among the leading causes of cancer death worldwide [1]. Systemic treatment has moved from tyrosine kinase inhibitors towards immune checkpoint inhibitor combinations such as atezolizumab plus bevacizumab [2], yet only a minority of patients respond, and markers that identify responders remain limited.

Ferroptosis is an iron-dependent form of regulated cell death driven by lipid peroxidation [3]. Glutathione peroxidase 4 (*GPX4*) is its central suppressor: it reduces phospholipid hydroperoxides to lipid alcohols, and loss of *GPX4* activity triggers ferroptosis in sensitive cancer cells [4]. In HCC cell lines, sorafenib has been reported to induce iron-dependent cell death [5], and activation of the p62-KEAP1-NRF2 pathway protects HCC cells from ferroptosis [6]. *GPX4* and the NRF2 antioxidant pathway are therefore both positioned as determinants of ferroptosis sensitivity in HCC.

Ferroptosis also interacts with anti-tumor immunity. CD8+ T cells release interferon-γ, which represses the cystine transporter *SLC7A11* and promotes tumor-cell ferroptosis [7], and ferroptosis of tumor-infiltrating neutrophils has been shown to be immunosuppressive [8]. These observations suggest that a ferroptosis-resistant, *GPX4*-high state could be accompanied by distinct immune and stromal features, but how *GPX4* expression relates to checkpoint expression and to immune and stromal composition in patient HCC has not been systematically examined.

NRF2 (encoded by *NFE2L2*) is a master regulator of the antioxidant response and has been reported to control ferroptosis-protective genes, including *GPX4* [9]. If *GPX4* expression in HCC were driven mainly by NRF2, *GPX4* would be expected to track NRF2 pathway activity. In addition, *CTNNB1* mutations, which are frequent in HCC, are associated with immune exclusion [10] and also reshape hepatocyte metabolism, so any association between *GPX4* and the tumor microenvironment needs to be examined with genotype and tumor purity in mind.

Here we used the TCGA-LIHC cohort to ask five questions: (i) is *GPX4* expression associated with immune checkpoints and with immune and stromal composition; (ii) do such associations survive adjustment for tumor purity, *CTNNB1* and *TP53* mutation, age and sex; (iii) which transcriptional programs distinguish *GPX4*-high from *GPX4*-low tumors; (iv) does *GPX4* track NRF2 activity; and (v) is *GPX4* expression prognostic. We report positive and null findings together, with effect sizes and confidence intervals, so that the data can be read for what they exclude as well as for what they show.

## MATERIALS AND METHODS

### 2.1 Data sources and cohort

RNA-sequencing data (RNA-Seq V2, RSEM-normalized gene expression), clinical data and somatic mutation calls for TCGA-LIHC [11] were downloaded from cBioPortal [12, 13] (study “Liver Hepatocellular Carcinoma, TCGA PanCancer Atlas”, lihc_tcga_pan_can_atlas_2018). Overall-survival data follow the TCGA Pan-Cancer Clinical Data Resource [14]. We retained primary tumor samples (TCGA sample type 01), one per patient, giving 366 tumors. Genes with missing or duplicated symbols were removed (20,511 genes retained) and expression was transformed as log_2_(RSEM + 1). Cohort characteristics are given in Table 1.

**Table 1.** Characteristics of the TCGA-LIHC cohort (PanCancer Atlas).

| **Characteristic** | **Value** |
|---|---|
| Primary tumors with RNA-seq, n | 366 |
| Age at diagnosis, years: median (IQR) | 61 (52-69) |
| Sex: female / male, n (%) | 120 (32.8%) / 246 (67.2%) |
| AJCC stage I / II / III / IV, n (% of 345 staged) | 169 (49.0%) / 86 (24.9%) / 84 (24.3%) / 6 (1.7%) |
| Histologic grade G1-2 / G3-4, n (% of 361 graded) | 227 (62.9%) / 134 (37.1%) |
| CTNNB1 non-synonymous mutation, n (% of 353 sequenced) | 94 (26.6%) |
| TP53 non-synonymous mutation, n (% of 353 sequenced) | 106 (30.0%) |
| KEAP1, NFE2L2 or CUL3 mutation, n (% of 353 sequenced) | 29 (8.2%) |
| Consensus tumor purity (CPE), median | 0.79 |
| GPX4 expression, log2(RSEM + 1): median (IQR); SD | 12.88 (12.38-13.35); 0.72 |
| Patients with overall-survival data, n (deaths) | 365 (129) |
| Median overall survival, months (Kaplan-Meier) | 55.4 |
| Median follow-up, months (reverse Kaplan-Meier) | 26.9 |

IQR, interquartile range. Percentages use the number of tumors with available data as denominator. Stage is the AJCC pathologic stage with sub-stages merged. Mutation status is based on non-synonymous coding variants.

### 2.2 Tumor purity and genotype

Consensus tumor-purity estimates (CPE) were taken from the supplementary data of Aran et al. [15] and matched to all 366 tumors by TCGA sample barcode. As a sensitivity check, a purity proxy based on somatic variant allele frequencies (twice the median variant allele frequency) was also used; it agreed only moderately with CPE (Spearman ρ = 0.48), and adjusted correlations obtained with either estimate differed by at most 0.04. Tumors were classed as mutant for a gene if they carried at least one non-synonymous coding variant (missense, nonsense, frame-shift, splice-site, in-frame insertion or deletion, translation-start or non-stop). Mutation status was analyzed for *CTNNB1*, *TP53* and, combined as an NRF2-pathway group, *KEAP1*, *NFE2L2* and *CUL3* (353 of 366 tumors had sequencing data).

### 2.3 Markers and gene-set scores

The primary panel comprised ten genes: *CD274* (PD-L1), *CD47*, *CD163*, *MRC1*, *NFE2L2*, *CTLA4*, *HAVCR2* (TIM-3), *LAG3*, *GZMA* and *PRF1*. The cytolytic activity score (CYT) was the geometric mean of *GZMA* and *PRF1* [16], computed as the mean of their log_2_ values, and *ACTA2* was analyzed as a myofibroblast marker. For the extended analysis we added single genes (*CD8A*, *CD3E*, *FOXP3*, *IFNG*, *CXCL9*, *PDCD1*, *TIGIT*) and gene-set scores: the 18-gene T-cell-inflamed gene-expression profile (GEP) of Ayers et al. [17]; and small curated marker sets for T cells (*CD3D*, *CD3E*, *CD3G*, *CD2*), CD8 T cells (*CD8A*, *CD8B*), NK cells (*NCR1*, *KLRC1*, *KLRD1*), B cells (*CD19*, *MS4A1*, *CD79A*, *CD79B*), macrophages (*CD68*, *CD14*, *CSF1R*), fibroblast/extracellular matrix (*COL1A1*, *COL1A2*, *COL3A1*, *DCN*, *LUM*), endothelium (*PECAM1*, *VWF*, *CDH5*, *CLDN5*) and NRF2 activity. The NRF2 activity score used 17 well-known NRF2 target genes (*NQO1*, *GCLM*, *GCLC*, *TXNRD1*, *SRXN1*, *HMOX1*, *AKR1B10*, *AKR1C1*, *AKR1C2*, *AKR1C3*, *SLC7A11*, *G6PD*, *PGD*, *TKT*, *ME1*, *OSGIN1*, *GSR*; *GPX4* was excluded); this list was curated by the authors and is not a published signature. A score is the mean of the per-gene z-scores of log_2_(RSEM + 1) across the cohort. All gene lists are given in Table S8. These marker-based scores are simple proxies for cell abundance and pathway activity, not validated deconvolution methods. Because the T-cell-inflamed profile contains *CD274* and *LAG3*, which were also analyzed individually, these results are not independent.

### 2.4 Correlation analysis

Associations with *GPX4* were estimated with Spearman’s rank correlation (ρ), which does not depend on the log transformation and is suited to skewed RSEM data. Confidence intervals used the Fisher transformation with the Bonett-Wright standard error, sqrt((1 + ρ^2^/2)/(n − 3)) [18] (in adjusted models, n − 3 is replaced by the residual degrees of freedom minus 1). Pearson correlations on log_2_ values are reported as a sensitivity analysis (Table S1). The ten-marker panel was corrected for multiple testing with the Benjamini-Hochberg false discovery rate (FDR) [19] within the panel (Table S1). In the extended analysis (Table S4), FDR was applied across all 28 targets for each model, and separately across four primary endpoints (T-cell-inflamed GEP, CD8 T-cell score, *CD274* and NRF2 activity score) that were fixed in an analysis plan (ANALYSIS_PLAN.md in the code repository) before the extended analyses were run on the TCGA data. The association with *CD274* had already been seen in the initial exploratory analysis, so it is not an independent test; all other associations should be regarded as exploratory. Tests were two-sided and FDR < 0.05 was taken as significant.

### 2.5 Adjusted analyses and gene-wide specificity

Adjusted associations were estimated as partial Spearman correlations: all variables were rank-transformed, *GPX4* and the target were regressed on the covariates, and ρ was derived from the t statistic of *GPX4* in the linear model (ρ = t/sqrt(t^2^ + df)). Three models were fitted: crude; adjusted for CPE purity; and adjusted for purity, *CTNNB1* mutation, *TP53* mutation, age (per 10 years) and sex (n = 352 tumors with complete data). Hepatitis etiology and vascular invasion were not available in the downloaded clinical files and could not be included. To judge whether *GPX4* is unusual among genes, we also correlated every expressed gene (mean log_2_(RSEM + 1) > 1; about 16,700 genes) with each of *CD274*, the T-cell-inflamed GEP, the CD8 T-cell score and the NRF2 activity score, and counted the proportion of genes whose correlation was at least as strong as that of *GPX4*.

### 2.6 Gene set enrichment analysis

Tumors were split at the median *GPX4* expression (183 high, 183 low). Genes were ranked by log_2_ fold change (mean log_2_(RSEM + 1) in *GPX4*-high minus *GPX4*-low tumors) and analyzed with gseGO in clusterProfiler [20] (Gene Ontology Biological Process, gene symbols, gene-set size 15-500, Benjamini-Hochberg adjusted p < 0.05, random seed 123), which uses the fgsea algorithm [21] of the GSEA method [22]. Because many highly significant terms share the smallest p-value the permutation procedure can return (adjusted p = 6.2 × 10^−9^), terms are ranked by normalized enrichment score (NES) rather than by p-value. The immune-related terms were counted by keyword search of term names (immune, leukocyte, T cell, lymphocyte, cytokine, interferon, antigen, phagocyt, macrophage, myeloid, inflammatory, B cell, natural killer, neutrophil, chemokine).

### 2.7 Survival analysis

Overall survival (OS) was analyzed in the 365 patients with survival data. Kaplan-Meier curves were compared between *GPX4*-high and *GPX4*-low tumors (median split) with the log-rank test. Cox proportional-hazards models [23] included log_2_ *GPX4* expression as a continuous variable (hazard ratio per 1 SD, SD = 0.72 log_2_ units, and per log_2_ unit) alone and with AJCC stage (I-IV), then additionally with age and sex, and with histologic grade (G1-2 versus G3-4). Proportional hazards were checked with Schoenfeld residuals (cox.zph); the global test was non-significant in all models (p = 0.21-0.31). Patients with missing stage were excluded from adjusted models (n = 344, 117 events).

### 2.8 Software and reproducibility

Analyses were run in R 4.4.2 (survival 3.7-0, survminer 0.5.1, clusterProfiler 4.14.6, fgsea 1.32.4, org.Hs.eg.db 3.20.0, ggplot2 4.0.1, patchwork 1.3.2); the extended analyses (adjusted correlations, gene-wide specificity, NRF2 context, extended Cox models) were run in R 4.3.3. Session information is recorded in the repository. All scripts, the analysis plan and instructions for downloading the input data are publicly available (see Code Availability).

## RESULTS

### 3.1 Cohort and *GPX4* expression

We analyzed 366 primary HCC tumors (Table 1). *GPX4* was expressed in all tumors (median log_2_(RSEM + 1) 12.88, interquartile range 12.38-13.35, SD 0.72). *GPX4* expression was not correlated with consensus tumor purity (ρ = 0.07, p = 0.16), whereas purity was strongly inversely correlated with immune and stromal scores in this cohort (T-cell-inflamed GEP ρ = −0.65; fibroblast/ECM score ρ = −0.43), so purity is a plausible confounder to be examined but was not simply collinear with *GPX4*.

### 3.2 *GPX4*-high tumors express less PD-L1 and *CD47*

*GPX4* expression was inversely correlated with PD-L1 (*CD274*; ρ = −0.34, 95% CI −0.43 to −0.24, p = 2.4 × 10^−11^, FDR = 2.4 × 10^−10^; Figure 1A) and with *CD47* (ρ = −0.21, 95% CI −0.31 to −0.11, p = 3.6 × 10^−5^, FDR = 1.2 × 10^−4^; Figure 1B; Table S1). Pearson correlations on log_2_ values had the same direction (R = −0.33 for *CD274* and −0.18 for *CD47*). None of the other checkpoint or cytotoxic markers in the panel was significant after correction: *CTLA4* ρ = −0.002, *LAG3* ρ = 0.04, *GZMA* ρ = 0.05, *PRF1* ρ = 0.01, and *HAVCR2* ρ = −0.11 (p = 0.031, FDR = 0.061).

**Figure 1.**
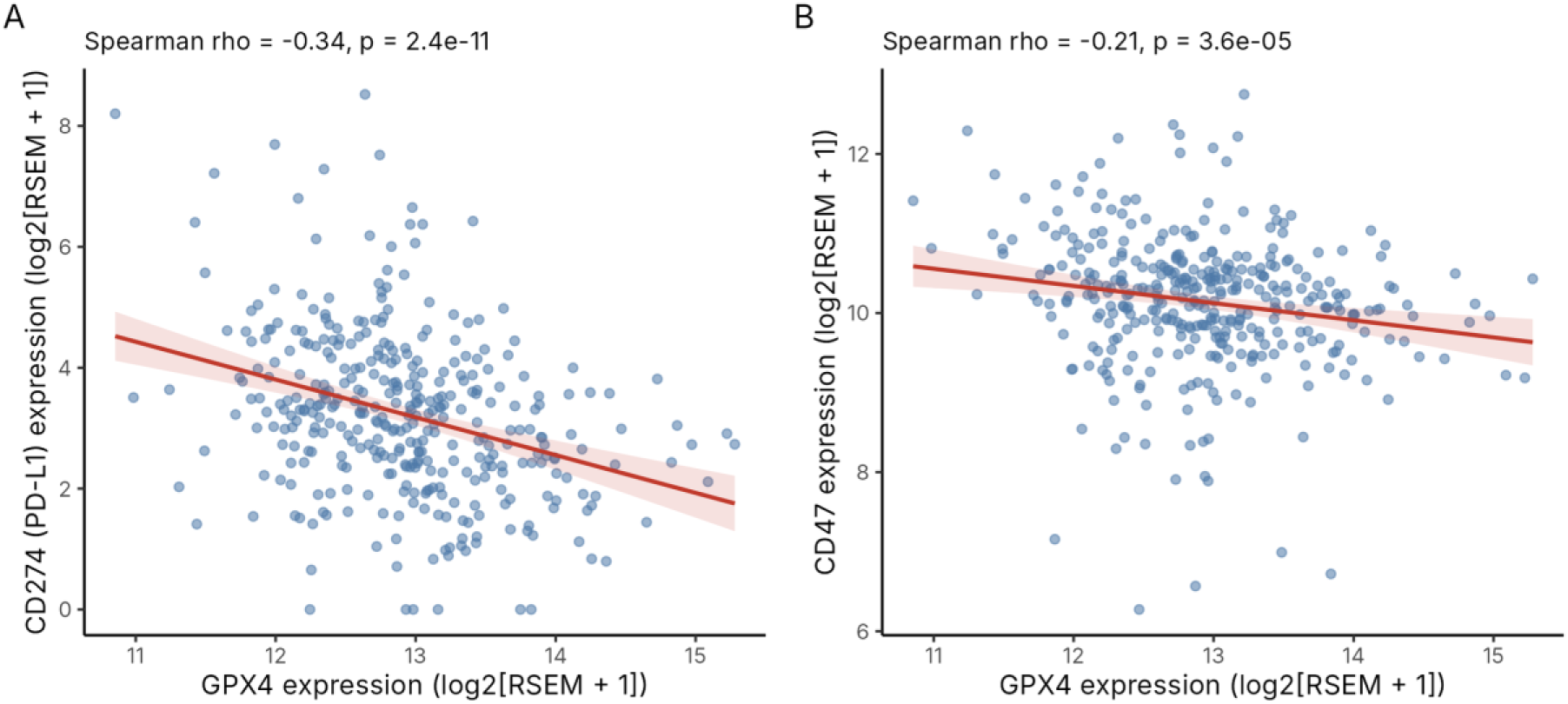
*GPX4* expression is inversely correlated with PD-L1 and *CD47*. **(A)** *GPX4* versus *CD274* (PD-L1) (Spearman ρ = −0.34, p = 2.4 × 10^−11^). **(B)** *GPX4* versus *CD47* (ρ = −0.21, p = 3.6 × 10^−5^). Each point is one primary tumor (n = 366); axes show log_2_(RSEM + 1). Lines are least-squares fits with 95% confidence bands for display; the statistics are rank-based.

### 3.3 Lower *CD163* and matrix-gene expression, but no association with T-cell or cytolytic scores

Among macrophage and stromal markers, *CD163*, a marker of M2-polarized macrophages, was inversely correlated with *GPX4* (ρ = −0.17, 95% CI −0.26 to −0.06, p = 0.0014, FDR = 0.0036; Figure 2A), whereas the second M2 marker *MRC1* was not (ρ = −0.03, p = 0.63; Figure 2B). A fibroblast/extracellular-matrix score (*COL1A1*, *COL1A2*, *COL3A1*, *DCN*, *LUM*) was lower in *GPX4*-high tumors (ρ = −0.22, 95% CI −0.32 to −0.12, p = 1.9 × 10^−5^), but the myofibroblast marker *ACTA2* was not (ρ = 0.10, p = 0.062; Figure 2C), so the association concerns collagen and matrix genes rather than all stromal-cell markers.

**Figure 2.**
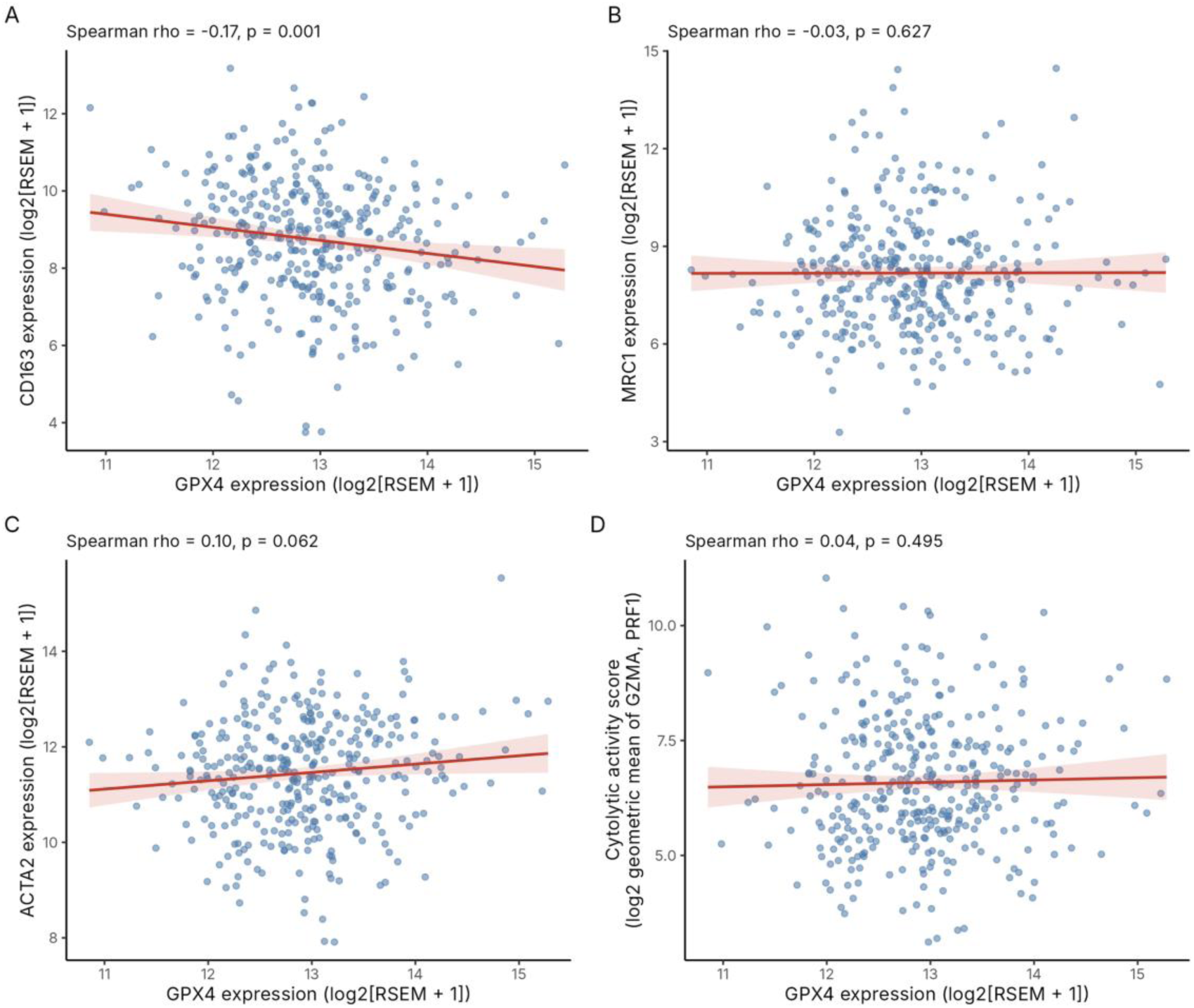
*GPX4* expression and macrophage, stromal and cytolytic markers. **(A)** *CD163* (ρ = −0.17, p = 0.0014). **(B)** *MRC1* (ρ = −0.03, p = 0.63). **(C)** *ACTA2* (ρ = 0.10, p = 0.062). **(D)** Cytolytic activity score, the geometric mean of *GZMA* and *PRF1* (mean of their log_2_ values; ρ = 0.04, p = 0.49). n = 366; axes show log_2_(RSEM + 1).

Markers of T-cell abundance and activity showed no association with *GPX4*. The cytolytic activity score (ρ = 0.04, 95% CI −0.07 to 0.14, p = 0.49; Figure 2D), the T-cell-inflamed GEP (ρ = −0.07, 95% CI −0.17 to 0.03, p = 0.17) and the CD8 T-cell score (ρ = −0.03, 95% CI −0.13 to 0.08, p = 0.62) all had confidence intervals that exclude correlations stronger than about 0.17 in either direction. T-cell, NK-cell and B-cell scores, and the individual genes *CD8A*, *CD3E*, *FOXP3*, *IFNG*, *CXCL9*, *PDCD1* and *TIGIT*, were also not significant after correction across all 28 targets (all FDR ≥ 0.07; Table S4). The data therefore do not indicate that *GPX4*-high tumors are depleted of, or enriched for, T-cell infiltration.

### 3.4 The associations persist after adjustment for purity and genotype

Adjusting for CPE purity left the main associations essentially unchanged (*CD274* ρ = −0.35, 95% CI −0.44 to −0.25; *CD47* ρ = −0.20; *CD163* ρ = −0.15; fibroblast/ECM score ρ = −0.21; Table 2, Figure 3). In the fully adjusted model (purity, *CTNNB1* and *TP53* mutation, age and sex; n = 352), *CD274* remained inversely associated with *GPX4* (ρ = −0.37, 95% CI −0.46 to −0.27, p = 8.0 × 10^−13^), as did *CD47* (ρ = −0.19, 95% CI −0.29 to −0.08, p = 4.4 × 10^−4^), *CD163* (ρ = −0.18, p = 6.5 × 10^−4^) and the fibroblast/ECM score (ρ = −0.17, p = 0.0011). The T-cell-inflamed GEP, the CD8 T-cell score and the NRF2 activity score remained null (Table 2). Among the four primary endpoints, only *CD274* was significant (FDR across the four endpoints 9.6 × 10^−11^ in the crude model).

**Figure 3.**
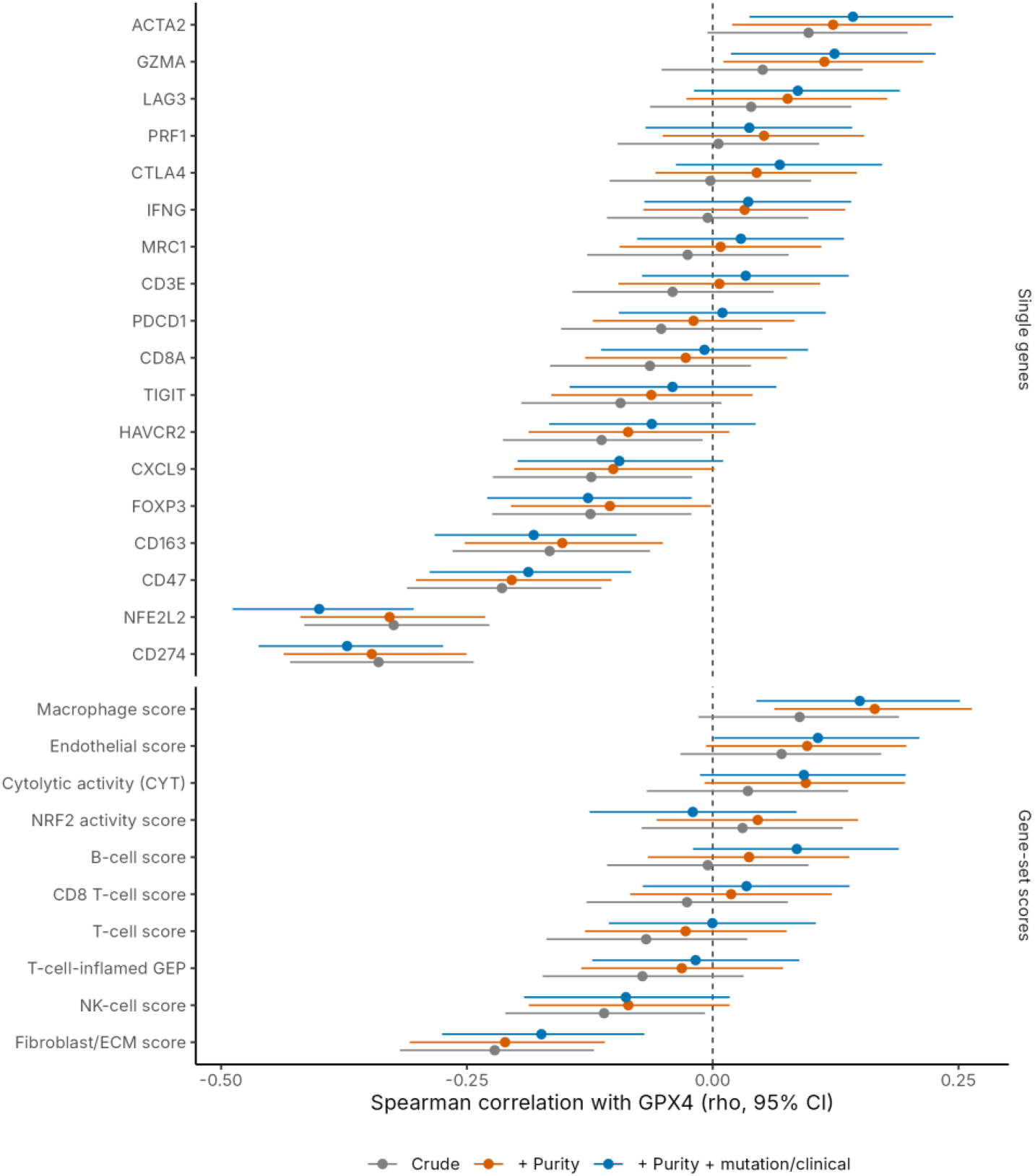
Association of *GPX4* with immune, stromal and NRF2 readouts before and after adjustment. Spearman ρ with 95% confidence intervals for single genes (upper panel) and gene-set scores (lower panel) in three models: crude (n = 366); adjusted for consensus tumor purity (n = 366); and adjusted for purity, *CTNNB1* and *TP53* mutation, age and sex (n = 352). Estimates for key targets are given in Table 2 and all estimates in Table S4. CYT, cytolytic activity score; GEP, gene-expression profile.

**Table 2.** Spearman correlation of *GPX4* expression with immune, stromal and NRF2 readouts, crude and adjusted.

| **Target** | **Crude ρ (95% CI)** | **+ Purity ρ (95% CI)** | **Fully adjusted ρ (95% CI)** | **p (fully adjusted)** |
|---|---|---|---|---|
| PD-L1 (CD274)† | −0.34 (−0.43, −0.24) | −0.35 (−0.44, −0.25) | −0.37 (−0.46, −0.27) | 8.0 × 10<sup>−13</sup> |
| CD47 | −0.21 (−0.31, −0.11) | −0.20 (−0.30, −0.10) | −0.19 (−0.29, −0.08) | 4.4 × 10<sup>−4</sup> |
| CD163 | −0.17 (−0.26, −0.06) | −0.15 (−0.25, −0.05) | −0.18 (−0.28, −0.08) | 6.5 × 10<sup>−4</sup> |
| Fibroblast/ECM score | −0.22 (−0.32, −0.12) | −0.21 (−0.31, −0.11) | −0.17 (−0.28, −0.07) | 0.0011 |
| NFE2L2 mRNA | −0.32 (−0.42, −0.23) | −0.33 (−0.42, −0.23) | −0.40 (−0.49, −0.30) | 8.8 × 10<sup>−15</sup> |
| ACTA2 | 0.10 (−0.01, 0.20) | 0.12 (0.02, 0.22) | 0.14 (0.04, 0.24) | 0.0078 |
| Macrophage score (CD68, CD14, CSF1R) | 0.09 (−0.01, 0.19) | 0.16 (0.06, 0.26) | 0.15 (0.04, 0.25) | 0.0052 |
| T-cell-inflamed GEP (18 genes)† | −0.07 (−0.17, 0.03) | −0.03 (−0.13, 0.07) | −0.02 (−0.12, 0.09) | 0.75 |
| CD8 T-cell score (CD8A, CD8B)† | −0.03 (−0.13, 0.08) | 0.02 (−0.08, 0.12) | 0.03 (−0.07, 0.14) | 0.52 |
| Cytolytic activity (CYT) | 0.04 (−0.07, 0.14) | 0.09 (−0.01, 0.20) | 0.09 (−0.01, 0.20) | 0.085 |
| NRF2 activity score (17 genes)† | 0.03 (−0.07, 0.13) | 0.05 (−0.06, 0.15) | −0.02 (−0.13, 0.09) | 0.71 |

Values are Spearman ρ with 95% confidence intervals (Bonett-Wright). “+ Purity”: adjusted for consensus purity estimate (n = 366). “Fully adjusted”: adjusted for purity, *CTNNB1* mutation, *TP53* mutation, age and sex (n = 352); the p-value refers to this model. †Primary endpoints defined in the analysis plan. ECM, extracellular matrix; GEP, gene-expression profile. False discovery rates and all other targets are given in Table S4.

Two results changed with adjustment. The three-gene macrophage score (*CD68*, *CD14*, *CSF1R*), which was not associated with *GPX4* in the crude analysis (ρ = 0.09, p = 0.091), became positively associated after purity adjustment (ρ = 0.16, 95% CI 0.06 to 0.26, p = 0.0016) and in the full model (ρ = 0.15, p = 0.0052), and *ACTA2* became positively associated in the full model (ρ = 0.14, p = 0.0078). Because *CD163* moves in the opposite direction to the macrophage score, the macrophage result depends on the marker used, and we do not conclude that *GPX4*-high tumors are macrophage-depleted. These macrophage and *ACTA2* analyses were not among the primary endpoints.

*GPX4* was not an outlier for the immune and NRF2 targets, but was in the tail for PD-L1. Of 16,722 expressed genes, 7.9% had an absolute correlation with *CD274* at least as large as that of *GPX4* (3.8% after purity adjustment); 1.3% were as strongly or more strongly negative (1.2% after adjustment; Figure S1, Table S7). For the T-cell-inflamed GEP, the CD8 T-cell score and the NRF2 activity score, 62.8%, 85.7% and 83.7% of genes, respectively, had an absolute correlation at least as large as that of *GPX4*, i.e. *GPX4* is unremarkable for these readouts.

### 3.5 *GPX4*-high tumors show metabolic enrichment and depletion of matrix and TGF-β programs

GSEA of *GPX4*-high versus *GPX4*-low tumors identified 1,459 significant GO Biological Process terms (291 with positive and 1,168 with negative NES; Table S2; Figure 4A shows representative terms). The strongest positive enrichments were respiration and oxidative-phosphorylation terms (aerobic electron transport chain, NES = 3.21; oxidative phosphorylation, NES = 3.14; all of the 12 highest-ranked terms had NES > 3.0), followed by mitochondrial translation (NES = 2.91). Ribosome biogenesis (NES = 2.15) and hepatocyte-type metabolic processes were also enriched, including xenobiotic metabolism (NES = 2.58), fatty-acid catabolism (NES = 2.55), cholesterol metabolism (NES = 2.26), bile-acid metabolism (NES = 1.93) and glutathione metabolism (NES = 1.88, adjusted p = 0.0028).

**Figure 4.**
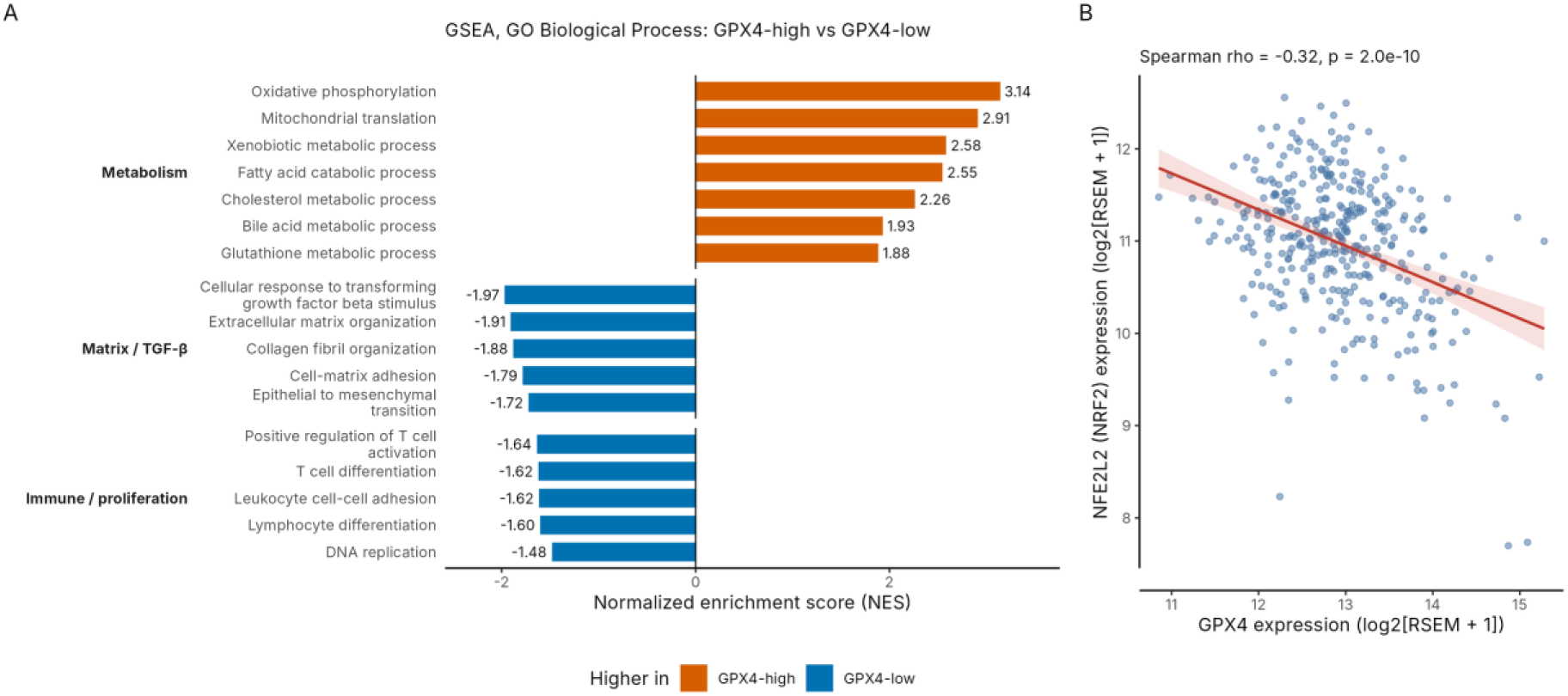
Transcriptional programs associated with *GPX4* expression, and the relationship between *GPX4* and *NFE2L2*. **(A)** Representative GO Biological Process terms from gene set enrichment analysis of *GPX4*-high versus *GPX4*-low tumors (median split), shown as normalized enrichment score (NES). Orange bars: higher in *GPX4*-high tumors; blue bars: higher in *GPX4*-low tumors. All terms shown have a Benjamini-Hochberg adjusted p < 0.05; the terms were chosen to illustrate the main themes and are not a random sample, and all 1,459 significant terms are listed in Table S2. **(B)** *GPX4* versus *NFE2L2* mRNA (ρ = −0.32, p = 2.0 × 10^−10^); axes show log_2_(RSEM + 1).

The most negative NES values were found for developmental and morphogenesis terms (glomerulus development, NES = −2.04; kidney development, NES = −1.95; Table S2). The matrix and TGF-β terms were also strongly negative: cellular response to TGF-β stimulus (NES = −1.97), extracellular-matrix organization (NES = −1.91), collagen fibril organization (NES = −1.88), cell-matrix adhesion (NES = −1.79) and epithelial-to-mesenchymal transition (NES = −1.72), consistent with the lower fibroblast/ECM score. Cell-cycle terms were also negatively enriched (all 24 terms containing “cell cycle”; DNA replication NES = −1.48). Of 130 significant terms whose names contained immune-related keywords, 125 had negative NES, including T-cell differentiation (NES = −1.62) and positive regulation of T-cell activation (NES = −1.64). These immune-labeled terms are large and overlap extensively with cell-adhesion, migration and developmental terms, and the targeted T-cell scores in Section 3.3 were null, so we do not regard the GSEA result as independent evidence of reduced T-cell presence.

### 3.6 *NFE2L2* mRNA is inversely correlated with *GPX4*, but NRF2 target-gene activity is not

*NFE2L2* mRNA was inversely correlated with *GPX4* (ρ = −0.32, 95% CI −0.42 to −0.23, p = 2.0 × 10^−10^; Figure 4B), and the association became stronger after adjustment (fully adjusted ρ = −0.40, 95% CI −0.49 to −0.30). It was present in tumors without *KEAP1*, *NFE2L2* or *CUL3* mutation (n = 324, ρ = −0.35) and in both *CTNNB1*-wild-type (n = 259, ρ = −0.36) and *CTNNB1*-mutant (n = 94, ρ = −0.42) tumors. However, *NFE2L2* mRNA is a poor readout of NRF2 activity, which is controlled mainly at the protein level by *KEAP1* [24]. In this cohort *NFE2L2* mRNA correlated only weakly with the 17-gene NRF2 activity score (ρ = 0.11, 95% CI 0.01 to 0.21, p = 0.037), and *GPX4* was not correlated with this score (ρ = 0.03, 95% CI −0.07 to 0.13, p = 0.56; Figure 5A, B; Table S5a). The estimate was similar in tumors without NRF2-pathway mutation (n = 324, ρ = 0.03), in *CTNNB1*-wild-type (ρ = −0.03) and *CTNNB1*-mutant tumors (ρ = 0.07), and in the 29 tumors with *KEAP1*, *NFE2L2* or *CUL3* mutation the estimate was imprecise (ρ = −0.16, 95% CI −0.50 to 0.23).

**Figure 5.**
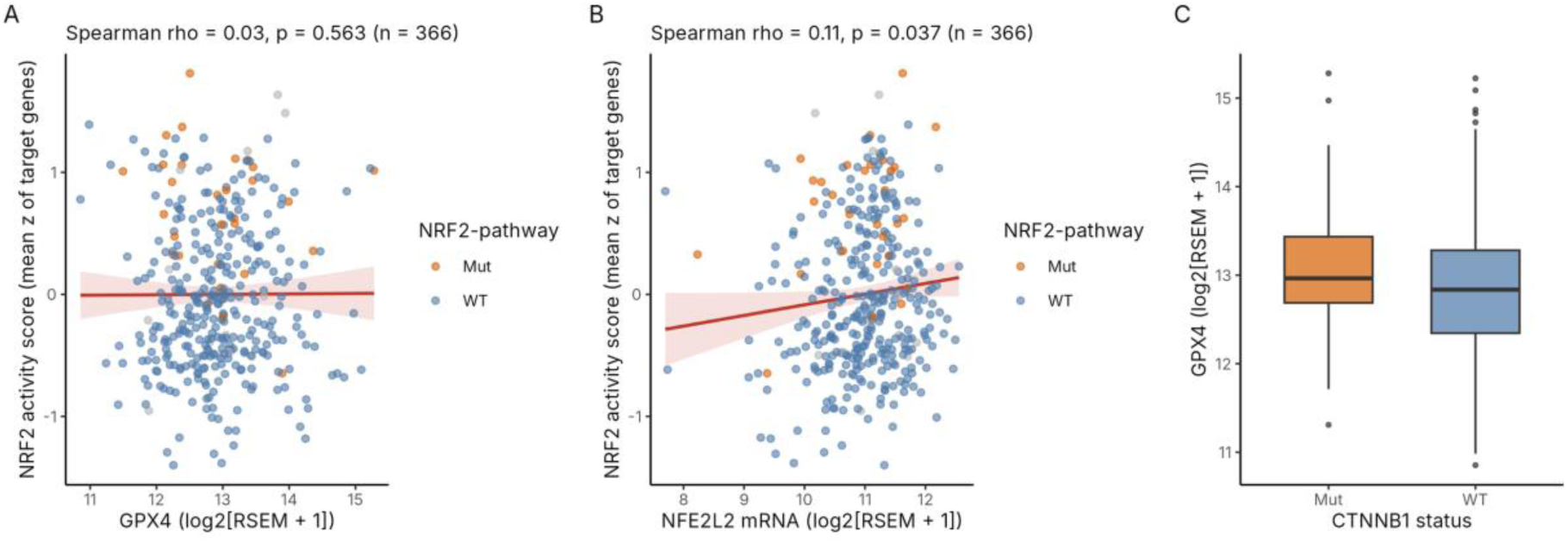
*GPX4*, *NFE2L2* and NRF2 target-gene activity. **(A)** *GPX4* versus the NRF2 activity score (mean z-score of 17 NRF2 target genes; ρ = 0.03, p = 0.56). **(B)** *NFE2L2* mRNA versus the same score (ρ = 0.11, p = 0.037). In A and B, orange points are tumors with a *KEAP1*, *NFE2L2* or *CUL3* mutation, blue points are wild type and grey points were not sequenced. **(C)** *GPX4* expression by *CTNNB1* mutation status (Wilcoxon p = 0.050); boxes show the median and interquartile range.

*GPX4* expression did not differ by *KEAP1*/*NFE2L2*/*CUL3* mutation (p = 0.94) or *TP53* mutation (p = 0.37) and was borderline higher in *CTNNB1*-mutant tumors (median 12.96 versus 12.84; Wilcoxon p = 0.050; Figure 5C). Individual NRF2 target genes were related to *GPX4* in opposite directions (Table S5c): *OSGIN1* (ρ = 0.38) and *AKR1C1*, *AKR1C2* and *AKR1C3* (ρ = 0.24 to 0.25) were positively correlated, whereas *GSR* (ρ = −0.42) and *GCLM* (ρ = −0.18) were negatively correlated, which explains why the composite score is flat.

### 3.7 *GPX4* expression is not associated with overall survival

Overall survival did not differ between *GPX4*-high (n = 182) and *GPX4*-low (n = 183) tumors (log-rank p = 0.25; Figure 6A; Tables S3 and S6). In Cox models, log_2_ *GPX4* expression was not associated with survival: hazard ratio per SD 1.03 (95% CI 0.87-1.22), p = 0.75 (365 patients, 129 deaths); adjusted for stage 1.09 (95% CI 0.91-1.29), p = 0.35 (n = 344, 117 events); additionally for age and sex 1.05 (95% CI 0.88-1.25), p = 0.61; additionally for grade 1.06 (95% CI 0.89-1.27), p = 0.52 (n = 342). Per log_2_ unit of *GPX4*, the stage-adjusted hazard ratio was 1.12 (95% CI 0.88-1.42). Comparing *GPX4*-high with *GPX4*-low tumors, the stage-adjusted hazard ratio was 1.34 (95% CI 0.93-1.94) (p = 0.12; Figure 6B). Stage was strongly prognostic in the same model (stage III versus I: HR 2.62, 95% CI 1.71-4.00; stage IV: HR 4.76, 95% CI 1.69-13.37). With 129 deaths and a median follow-up of 26.9 months, the data are compatible with no association and with a modest adverse effect (upper confidence limits 1.22-1.29 per SD), but not with a large one.

**Figure 6.**
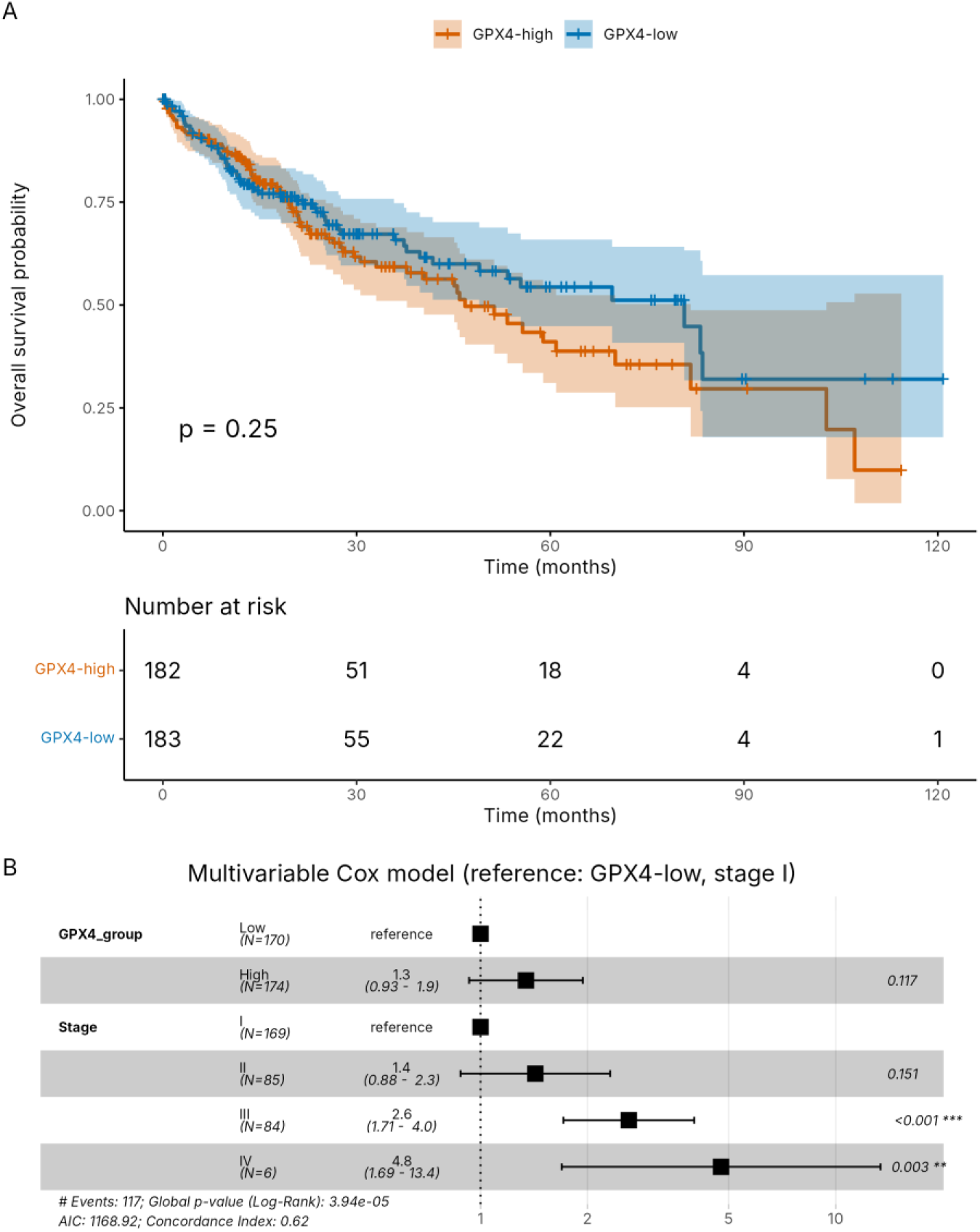
*GPX4* expression and overall survival. **(A)** Kaplan-Meier overall survival for *GPX4*-high (n = 182) and *GPX4*-low (n = 183) tumors, split at the median (log-rank p = 0.25); shaded areas are 95% confidence intervals and numbers at risk are shown below the plot. **(B)** Multivariable Cox model with *GPX4* group and AJCC stage (n = 344, 117 events); reference categories are *GPX4*-low and stage I.

## DISCUSSION

### 4.1 Principal findings

In 366 primary HCC tumors, high *GPX4* expression marked a coherent transcriptional state: lower PD-L1 and *CD47* expression, lower collagen/extracellular-matrix and TGF-β programs, and higher oxidative-phosphorylation, mitochondrial, lipid, xenobiotic and glutathione metabolism. These associations were moderate in size (|ρ| 0.17-0.34, i.e. at most 12% of rank variance) but robust to adjustment for tumor purity, *CTNNB1* and *TP53* mutation, age and sex. Equally informative are the null results: *GPX4* was not associated with T-cell-inflamed, CD8 T-cell or cytolytic scores, with an NRF2 target-gene score, or with survival.

### 4.2 A checkpoint-low state that is not a T-cell-excluded state

Lower PD-L1 and *CD47* expression in *GPX4*-high tumors could be read as a “cold”, immune-excluded phenotype. Our data do not support that reading. The T-cell-inflamed profile, CD8 score and cytolytic score were unrelated to *GPX4*, with confidence intervals that exclude all but weak associations, and the single-gene T-cell markers were also null. The checkpoint association is therefore better described as a lower expression of PD-L1 and *CD47* transcripts at similar T-cell content. Because PD-L1 and *CD47* mRNA in bulk tumor tissue come from tumor cells and from stromal and myeloid cells, one possible explanation is that *GPX4*-high tumors contain fewer matrix-producing and PD-L1-expressing non-tumor cells, which would also explain the parallel decline of *CD163* and of the fibroblast/ECM score. The persistence of the associations after CPE adjustment argues against simple dilution by non-tumor cells, but CPE is an imperfect estimate of stromal content, and we did not test cell-type origin directly (for example by single-cell data). *CD47* has been proposed as an immunotherapy target in HCC [25]; whether lower *CD47* mRNA in *GPX4*-high tumors has any therapeutic meaning is unknown.

### 4.3 A metabolically active, matrix-poor state

GSEA, which is unbiased by our choice of markers, points to a coherent metabolic program in *GPX4*-high tumors (oxidative phosphorylation, mitochondrial translation, fatty-acid catabolism, xenobiotic, cholesterol, bile-acid and glutathione metabolism) together with reduced TGF-β, collagen and cell-adhesion signatures. Many of these are hepatocyte functions, so *GPX4*-high tumors may be more hepatocyte-like or better differentiated, and oxidative, lipid-rich metabolism provides substrates for lipid peroxidation, which would make a high *GPX4* level necessary for survival. These are hypotheses: we did not test *GPX4* against histologic grade, molecular subtype or differentiation markers, and the enrichment results are correlative. We use “stroma-poor” in the title in this limited sense, referring to lower collagen/matrix and TGF-β programs; *ACTA2* and endothelial markers were not lower, so the stromal compartment is not uniformly reduced.

### 4.4 *GPX4* and NRF2

*GPX4* was inversely correlated with *NFE2L2* mRNA across strata, but not with NRF2 target-gene activity. *NFE2L2* mRNA correlated only weakly with the activity score, which supports the view that mRNA level is a poor proxy for NRF2 activity [24]. The inverse mRNA correlation is therefore not evidence that *GPX4* is uncoupled from NRF2 signaling, and the null result for the activity score is limited by the score itself: it is a curated set of 17 genes that were not validated against NRF2 perturbation in liver tumors, and several of its members, such as *GSR* and *GCLM*, are inversely correlated with *GPX4* whereas *OSGIN1* and the AKR1C genes are positively correlated. Protein-level or perturbation data would be needed to determine whether NRF2 regulates *GPX4* in HCC.

### 4.5 Prognosis and clinical relevance

*GPX4* expression was not prognostic in this cohort with or without adjustment for stage, age, sex or grade. We make no claim that *GPX4*-high status predicts response to immune checkpoint blockade: no treatment data were analyzed, the T-cell scores did not differ, and patients in the pivotal atezolizumab plus bevacizumab trial were not selected by PD-L1 expression [2]. The findings are hypothesis-generating for experiments that test whether *GPX4* dependence and tumor metabolism and matrix state are linked.

### 4.6 Limitations

This is a single-cohort, correlative analysis of bulk mRNA data. No independent validation cohort was analyzed, and the results should be confirmed in external HCC datasets with RNA-sequencing and clinical annotation. The immune and stromal scores are small marker sets (two to five genes for several of them) rather than validated deconvolution methods, and the NRF2 target list is not a published signature. Hepatitis etiology and vascular invasion were not available and were not adjusted for. Twenty-eight targets were tested in the extended analysis, only four were designated primary, and the *CD274* association was known before the plan was written; secondary findings (macrophage score, *ACTA2*, GSEA themes) are exploratory. Only six patients had stage IV disease and median follow-up was 26.9 months, which limits the survival analysis. Finally, mRNA levels do not measure protein abundance, enzymatic activity or ferroptosis sensitivity, and no functional experiments were performed.

## CONCLUSIONS

High *GPX4* expression in hepatocellular carcinoma marks a metabolically active tumor state with lower PD-L1, *CD47* and matrix/TGF-β program expression. This state is not accompanied by differences in T-cell-inflamed, CD8 or cytolytic scores, by NRF2 target-gene activity, or by survival. The observations are hypothesis-generating and require external validation and functional testing.

## DATA AVAILABILITY STATEMENT

The RNA-sequencing, clinical and mutation data analyzed in this study are publicly available from the TCGA-LIHC PanCancer Atlas study on cBioPortal (https://www.cbioportal.org; study ID lihc_tcga_pan_can_atlas_2018). Consensus tumor-purity estimates are available in the supplementary data of Aran et al. (2015). No new data were generated. Supplementary Tables S1-S8 are provided with this preprint.

## CODE AVAILABILITY STATEMENT

The analysis scripts (Master_Analysis_Script.R and Extended_Analysis_Script.R), the pre-analysis plan (ANALYSIS_PLAN.md) and a README with instructions to reproduce all figures and tables are publicly available at https://github.com/prakas000/GPX4-HCC-Immune-Metabolism. Session information for each run is written to the output folder by the scripts.

## Supporting information

Supplementary Tables S1-S8 (Excel workbook)

## SUPPLEMENTARY MATERIAL

Supplementary Tables S1-S8 are provided as a separate Excel workbook (Supplementary_Tables_S1-S8.xlsx):

- Table S1. Spearman and Pearson correlations of *GPX4* with the ten-marker panel, the cytolytic activity score and *ACTA2* (n = 366).
- Table S2. Significant GO Biological Process terms from GSEA of *GPX4*-high versus *GPX4*-low tumors (1,459 terms).
- Table S3. Cox models of overall survival: continuous and median-split *GPX4*, alone and adjusted for stage.
- Table S4. Crude and adjusted correlations of *GPX4* with 28 immune, stromal and NRF2 targets (Spearman and Pearson), with FDR.
- Table S5. NRF2 context: (a) correlations by mutation stratum; (b) *GPX4* expression by mutation status; (c) correlations of *GPX4* with individual NRF2 target genes.
- Table S6. Extended Cox models (hazard ratio per 1 SD of log_2_ *GPX4*), with Schoenfeld tests.
- Table S7. Gene-wide specificity: (a) proportion of genes correlated with each target at least as strongly as *GPX4*; (b) top correlated genes.
- Table S8. Gene sets used for the signature scores.

**Figure S1.**
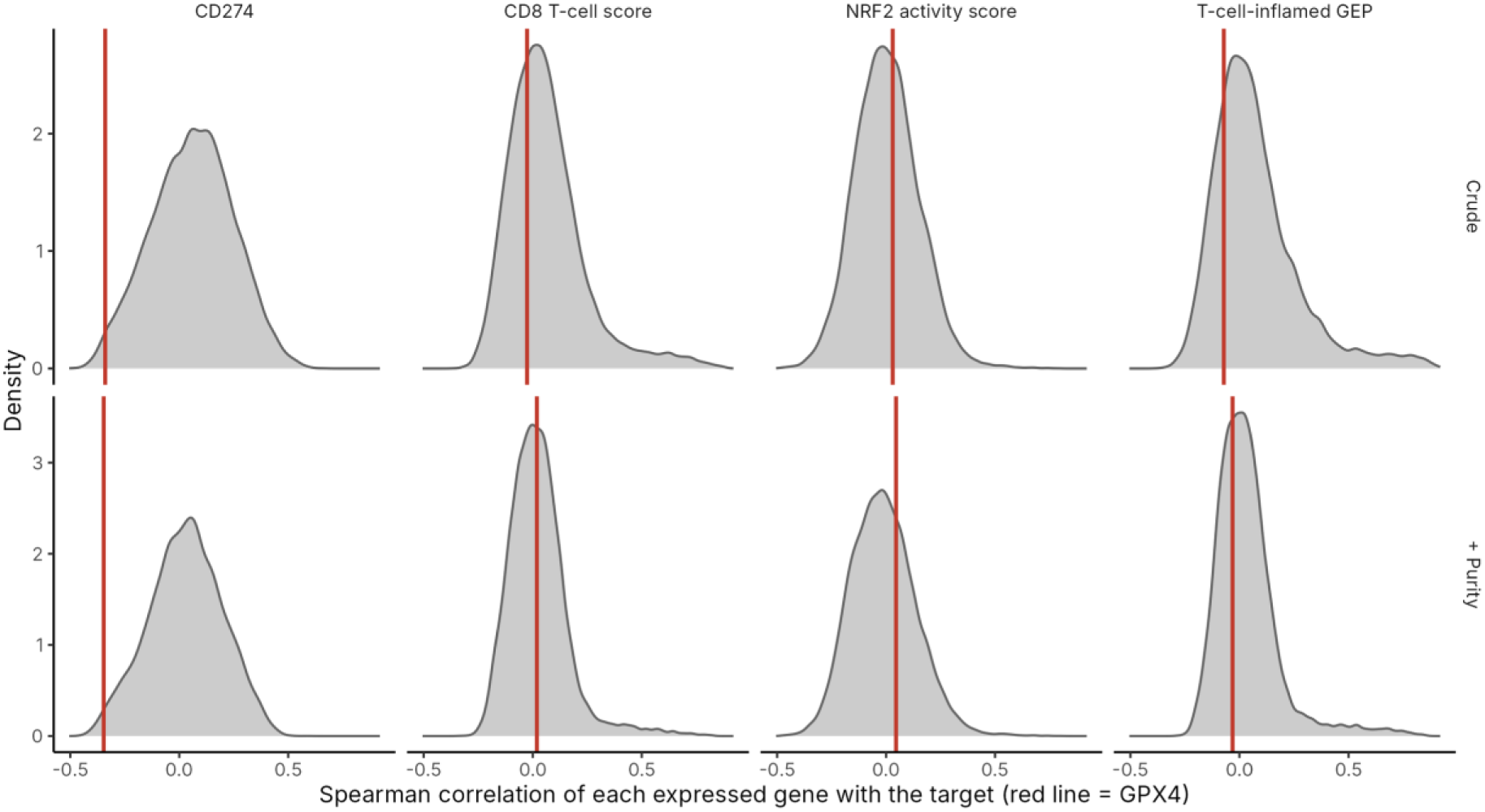
Gene-wide specificity of the *GPX4* correlations. Distribution of Spearman correlations between each expressed gene (mean log_2_(RSEM + 1) > 1) and *CD274*, the CD8 T-cell score, the NRF2 activity score and the T-cell-inflamed profile, without adjustment (upper row) and adjusted for purity (lower row). The red line marks the correlation of *GPX4* with the same target.

